# Single-cell transcriptome analyses recapitulate the cellular and developmental responses to abiotic stresses in rice

**DOI:** 10.1101/2020.01.30.926329

**Authors:** Yu Wang, Qing Huan, Xiao Chu, Ke Li, Wenfeng Qian

## Abstract

The metabolism and reproduction of plants depend on the division of labors among cells. However, cells with various functions are often studied as a bulk where their specificities were diluted. Here, we apply single-cell RNA sequencing to the aerial part of rice seedlings growing in various environments. We capture the transcriptomes of thousands of cells, identify all major cell types, and reconstruct their developmental trajectories. We find that abiotic stresses not only affect gene expression in a cell-type-specific manner but also have impacts on the physical size of cells and the composition of cell populations. We validate some of these conclusions with microscopy and provide developmental mechanisms with computational analyses. Collectively, our study represents a benchmark-setting data resource of single-cell transcriptome atlas in rice and an illustration of exploiting such resource to drive discoveries in plant biology.

## INTRODUCTION

Multicellular plants are composed of cells with a variety of types, which function as a whole to warrant the metabolism and reproduction of the organisms. Since somatic cells carry largely identical genetic information, their differences are likely created from the epigenetic regulation of gene expression. Therefore, understanding heterogeneity in gene expression among cells or across environments offers new insights into the development and physiology of plants. However, such heterogeneity is often missed in the traditional bulk RNA sequencing (RNA-seq) analyses since the gene expression pattern in a small fraction of cells can be diluted by that in others.

Individual plant cells or cell types have been isolated with glass microcapillaries (Han et al., 2017; Lieckfeldt et al., 2008), laser microdissection (Frank and Scanlon, 2015; Jiao et al., 2009), microscopic isolation of reporter-expressing protoplasts, and fluorescence-activated cell sorting (Birnbaum et al., 2003; Dinneny et al., 2008; Efroni et al., 2015). Combined with microarray or RNA-seq, these analyses have shown widespread heterogeneous and monoallelic gene expression, identified motifs in cell-type-specific promoters, etc. However, these approaches are often limited to cells with prior knowledge of locations in a plant or tissue-specific promoters.

The last decade has witnessed the development and application of single-cell RNA sequencing (scRNA-seq) techniques that can characterize cells of different types in a single assay. One of the strategies is to encapsulate cells into individual liquid droplets using a microfluidic device, and therefore, the transcriptomes of thousands of cells were obtained in a single experiment; identifies of individual cells are further recognized with following-up computational analyses (Klein et al., 2015; Macosko et al., 2015). Such strategy was mainly applied to animal tissues, although a growing number of applications in plants were implemented in the year of 2019 (Denyer et al., 2019; Jean-Baptiste et al., 2019; Rich-Griffin et al., 2019; Ryu et al., 2019; Shulse et al., 2019; Zhang et al., 2019). All these studies have been focused on the *Arabidopsis* root; the application of scRNA-seq to other plant cells remains absent, presumably because less prior knowledge on the cellular composition and developmental processes makes it technically more challenging.

In this study, we performed scRNA-seq in the aerial part of rice seedlings in various conditions, including facing two abiotic stresses. We explored cellular and developmental mechanisms in responding to stresses from the single-cell transcriptome data, by identifying cell-type-specific regulation of gene expression, detecting changes in the total number of transcripts in a cell, and investigating the compositional variation in cell populations. Our study showcases the power of the application of scRNA-seq on the aerial part of plants and paves the way for the construction of plant cell atlas (Rhee et al., 2019).

## RESULTS

### Single-cell mRNA sequencing identifies major cell types in the aerial part of rice seedlings

To study cellular heterogeneity during stress responses, we attempted to recognize individual cell types using single-cell mRNA sequencing (scRNA-seq). We isolated protoplasts from the aerial part of two-week-old rice seedlings growing in Kimura B nutrient solution (“control” hereafter), treated with 200 mM additional NaCl for three days (high-salinity), or treated with a low concentration of ammonium cation for five days (20 mM NH_4_^+^, low-nitrogen). We performed scRNA-seq on the 10x Genomics platform (Zheng et al., 2017) and Illumina sequencing (**Figure 1A**). We obtained transcriptomes for a total of 4580 cells, among which the medians of unique molecular identifiers (UMIs) and genes were 10256 and 2480 per cell, respectively. When sequencing reads were aggregated among individual cells (“pseudobulk” hereafter), we detected the expression of 25822 out of 39045 non-transposable element-related (non-TE-related) genes in the rice genome (66.4%, **Table S1**), a percentage similar to that in an accompanied bulk RNA-seq experiment when a cutoff of transcript per million (TPM) > 0.5 was applied (61.3%). Furthermore, the expression level estimated in bulk RNA-seq and in the pseudobulk were significantly correlated among genes (Spearman’s correlation coefficient *ρ* = 0.72, *P* < 10^−100^, **Figure S1**). Collectively, these observations indicate that the landscape of rice transcriptome was largely reproduced in our scRNA-seq experiments.

**Figure 1.**
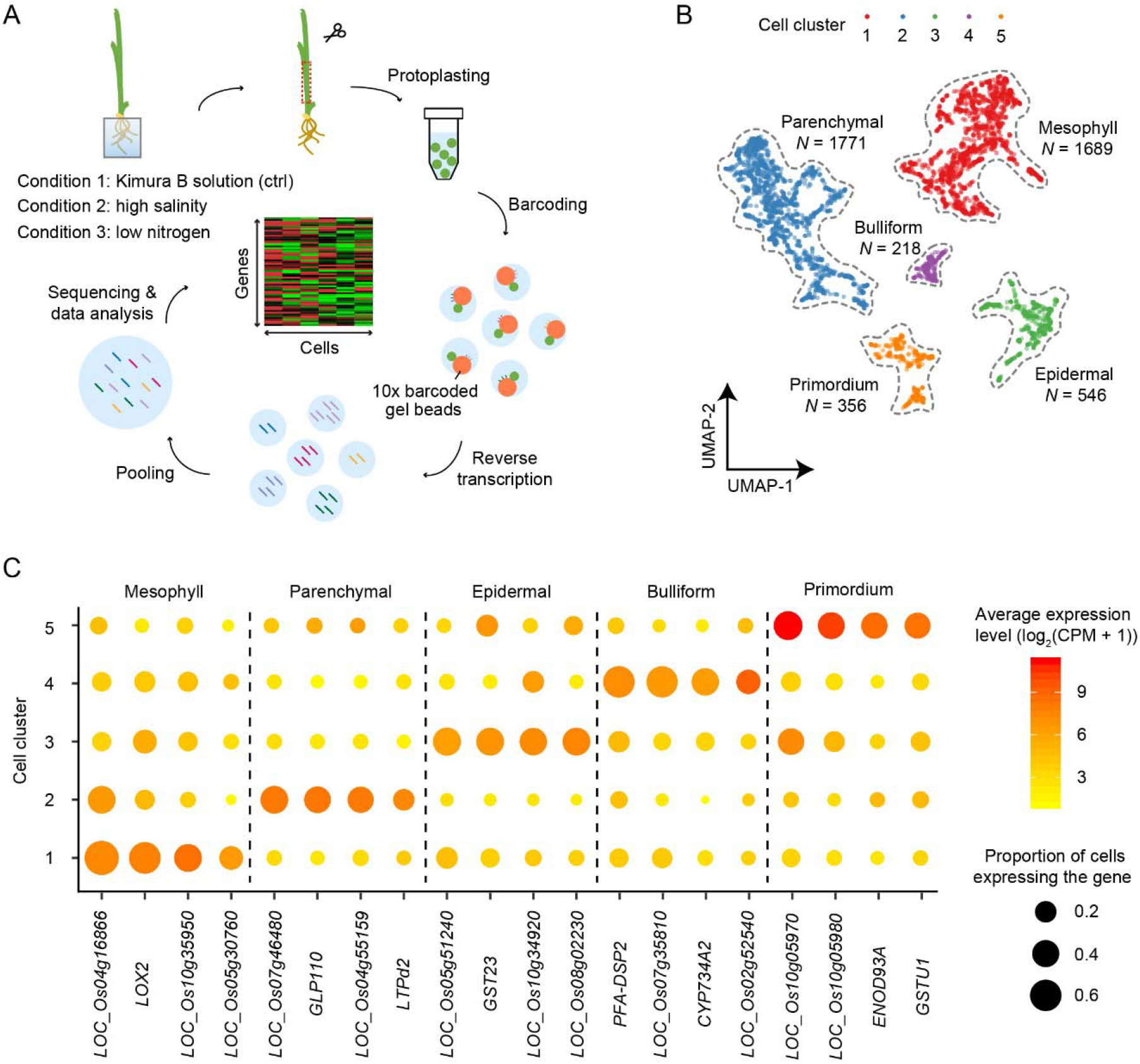
Single-cell transcriptome atlas of the areal part of rice seedlings. (A) A schematic diagram shows the workflow of scRNA-seq. (B) The dimension reduction by UMAP and the classification of five major cell clusters. Dots represent individual cells and are colored according to the cell clusters. The number of cells (*N*) in each cluster is shown. (C) The expression pattern of cluster-specific genes. The average expression level (color) and the proportion of cells expressing the gene (area) are shown for each gene and cell cluster. Genes are labeled with their common or systematic names. The detailed information of cluster-specific genes is provided in Table S3.

To recognize the type of cells detected in the scRNA-seq experiment, we clustered cells according to their transcriptomes. We excluded 63 genes that are differentially expressed during protoplasting (**Table S2**). We estimated gene expression for individual cells in the unit of count per million (CPM), which controls the total number of transcripts captured in a cell (or the total UMI of all genes in a cell). We identified 1620 most variable genes among cells and, based on the similarity in their expression, projected individual cells onto a two-dimensional space using uniform manifold approximation and projection (UMAP) (McInnes et al., 2018). To ensure coherent cell-type assignment among samples in various environments, we corrected for the batch effect by detecting the mutual nearest neighbors (MNN) (Haghverdi et al., 2018). We classified cells into five major clusters (McInnes et al., 2017), each containing a variable number of cells from 218 to 1771 (**Figure 1B**). We assigned cells to five types according to the cluster-specific expression of genes of known function — mesophyll, parenchymal, epidermal, bulliform, and primordium cells (**Figure 1C** and **Table S3**). Consequently, all major cell types in the aerial part of rice seedlings were identified in our scRNA-seq experiments.

### Abiotic-stress-induced transcriptomic changes vary among cell types

Not unexpectedly, the seedling samples of control, low-nitrogen, and high-salinity contained all five cell types (**Figure 2A**). To detect if the expression of some genes is affected by abiotic stresses in a cell-type-specific manner, we identified differentially expressed genes (DEGs) for each cell type upon the low-nitrogen or high-salinity treatment. We defined DEGs as genes with a ≥ 2-fold difference in the average expression level and *P* < 0.01 in the Mann-Whitney *U* test (two examples in **Figure 2B**, others in **Figure S2**). DEGs identified in the low-nitrogen and high-salinity treatments are shown in **Figure 2C** and **Figure S3A**, respectively. Heterogeneity in gene expression response during stress was observed among cell types; although shared gene expression regulation existed, the majority of genes response to stresses in a cell-type-specific manner. This is consistent with previous observations in Arabidopsis roots that the expression of 49% of DEGs was only altered in a single cell type upon sucrose removal (Shulse et al., 2019).

**Figure 2.**
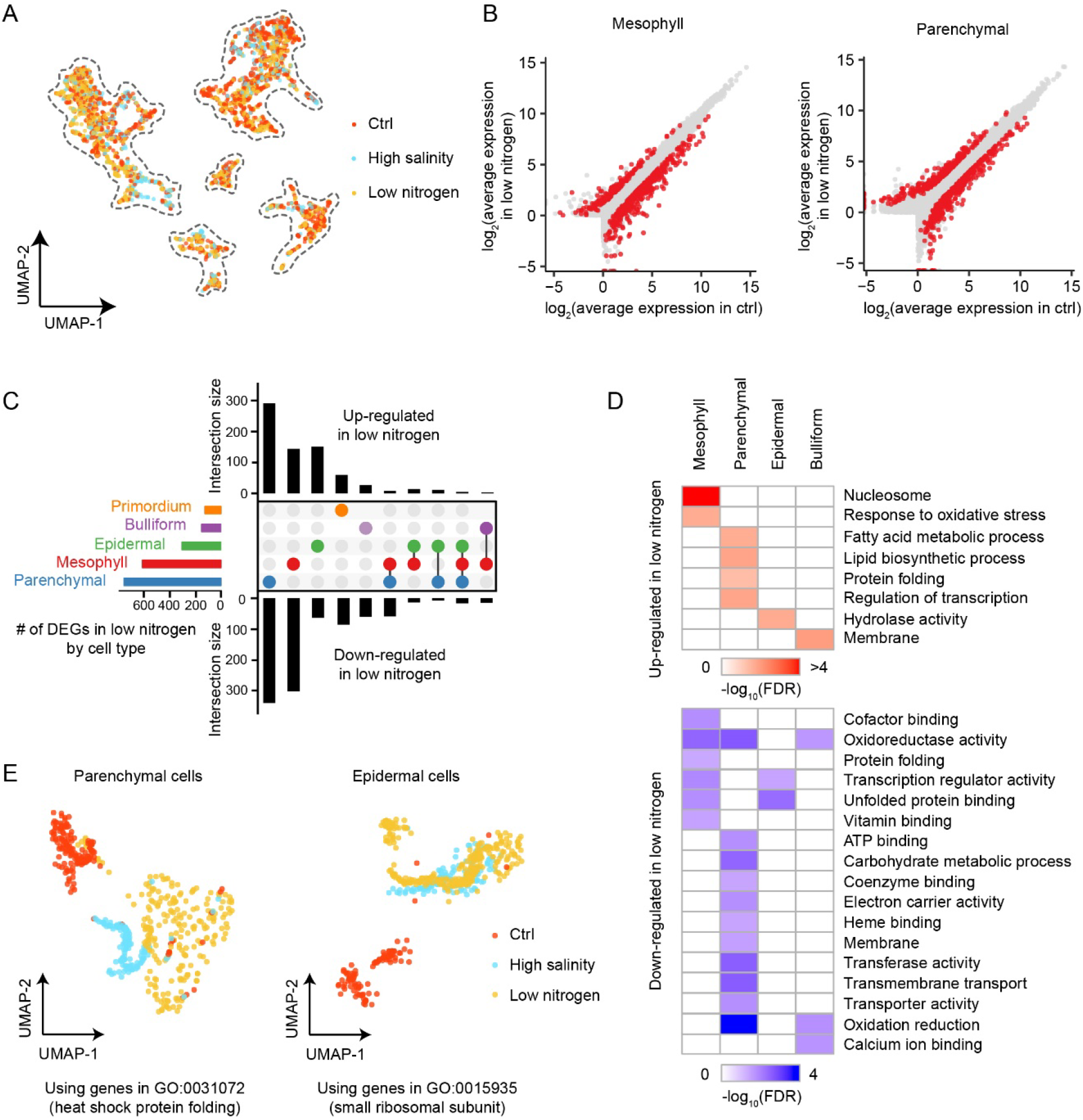
Cell-type-specific transcriptome responses upon abiotic stresses. (A) The three differently treated samples largely overlap each other in the UMAP plot. (B) Scatter plots show genes that are differently expressed (colored in red) under low-nitrogen treatment in mesophyll and parenchymal cells, respectively. The identification of DEGs in other conditions or cell types is shown in Figure S2. (C) Number of DEGs responding to the low-nitrogen stress that are cell-type-specific or shared by multiple cell types. The bar plot on the left shows the number of DEGs (including both up- and down-regulated) for each cell type. The upper and lower bar plots show the number of up- and down-regulated DEGs, respectively. The numbers of DEGs upon the high-salinity stress are shown in Figure S3A. (D) DEG-enriched GO terms responding to the low-nitrogen stress in each cell type. DEGs in primordium cells do not have enriched GO terms. DEG-enriched GO terms responding to the high-salinity stress are shown in Figure S3B. (E) When a subset of genes (all genes in a GO term) are used, cells treated with various abiotic stresses cluster together in the UMAP plot and separate from those from the control sample, suggesting that the transcriptional regulation of these genes is shared by the responses to various abiotic stresses.

DEGs identified in individual cell types of rice were enriched in various gene ontology (GO) terms (**Figure 2D** and **Figure S3B**). For example, in responding to low nitrogen, genes whose expression was induced in mesophyll cells were enriched in response to oxidative stress (false discovery rate, FDR = 0.02). Notably, genes whose expression was suppressed in the same cell type and in responding to the same stress were also enriched in oxidoreductase activity. These observations suggest an intrinsic coupling between oxidative and ammonium metabolisms in mesophyll cells.

### Common transcription programs are rewired in response to high salinity and low nitrogen

DEGs were significantly overlapped between the high-salinity and low-nitrogen treated samples in each of the five cell types (e.g., odds ratio = 3.6 and 2.9 for mesophyll and parenchymal cells, respectively, **Figure S3C**); that is, genes showing differential expression during the low-nitrogen treatment tended to be differentially expressed during the high-salinity treatment, implying that plants likely respond to various abiotic stresses with common rewiring strategies of the regulatory networks. Consistently, when we performed feature projection using only genes in a GO category, cells under high-salinity or low-nitrogen treatment clustered together and fairly separated from those in the control sample (two examples are shown in **Figure 2E**).

### Reduction in mRNA population captured in scRNA-seq reveals that epidermal cells became physically smaller during stress responses

The total UMI in a cell, which reflects the global mRNA population captured in an scRNA-seq experiment, significantly varied upon stresses in some cell types (**Figure 3A**). In particular, the total UMI in epidermal cells significantly decreased upon both low-nitrogen and high-salinity treatments. The total UMI in a cell has been reported to be positively correlated with its physical size in humans (Chen et al., 2019; Kempe et al., 2015); alternatively, the total UMI could simply reflect the activity of RNA polymerase in a cell. To distinguish these possibilities we estimated cell size from microscopic images. The areas of epidermal cells reduced upon both stresses, particularly in high salinity (**Figures 3B–C**), indicating that the total UMI per cell can be used to infer the size of epidermal cells.

**Figure 3.**
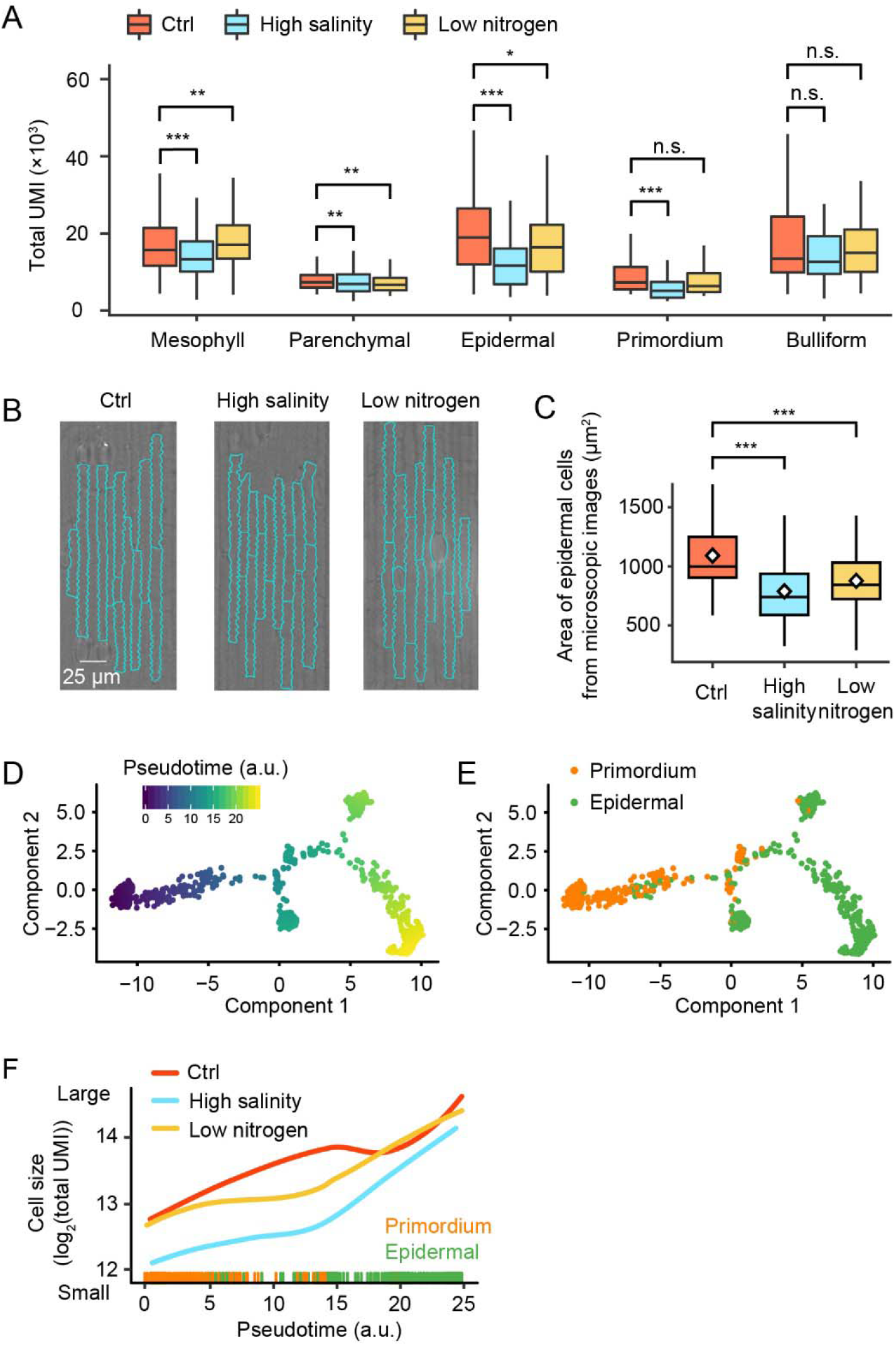
Cell-type-specific cell-size responses upon abiotic stresses. (A) The total UMI (i.e., the number of transcripts captured by scRNA-seq) significantly decreases upon the high-salinity and low-nitrogen stresses in some cell types. *P* values are given by the Student’s *t*-tests (significance level: ***, *P* < 0.001; **, *P* < 0.01; *, *P* < 0.05; n.s., not significant). (B–C) Cell sizes of epidermal cells in the control and the stress-treated samples. Statistical analyses are performed with 67, 59, and 70 cells, in ctrl, high-salinity, and low-nitrogen samples, respectively. *P* values are given by the Student’s *t*-tests (***, *P* < 0.001). (D–E) Developmental trajectory from primordium to epidermal cells. Each dot represents a cell and is colored by pseudotime (D) or cell types (E). (F) The cell size (as inferred from the total UMI in a cell) increases along the developmental pseudotime. The rate varies among three conditions.

To understand the underlying mechanisms that led to the reduced size of epidermal cells upon stresses, we reconstructed the developmental trajectory from primordium to epidermal cells (**Figures 3D–E**). While the total UMI increased over the development in all three conditions, the rates were different (**Figure 3F**). Plants growing in the low-nitrogen condition showed a delay in cell expansion during the primordium-epidermal cell transition. In addition to that, plants growing in the high-salinity condition showed an overall reduced size of both primordium and epidermal cells, possibly related to the increased osmotic pressure that leads to a reduced turgor pressure against cell wall. Collectively, these results reveal a cell-type-specific change in size responding to abiotic stresses.

### Abiotic stress decelerates the differentiation towards mesophyll cells

The composition of cell populations also changed upon abiotic-stress treatments (**Figure 4A**). In particular, the proportion of mesophyll cells decreased, and that of parenchymal cells increased in responding to either high salinity or low nitrogen. Consistently, under microscopes we observed a ~ 50% reduction in the intensity of red autofluorescence, which was emitted from chloroplasts that are mainly present in the mesophyll cells (**Figures 4B–C** and **Figure S4**).

**Figure 4.**
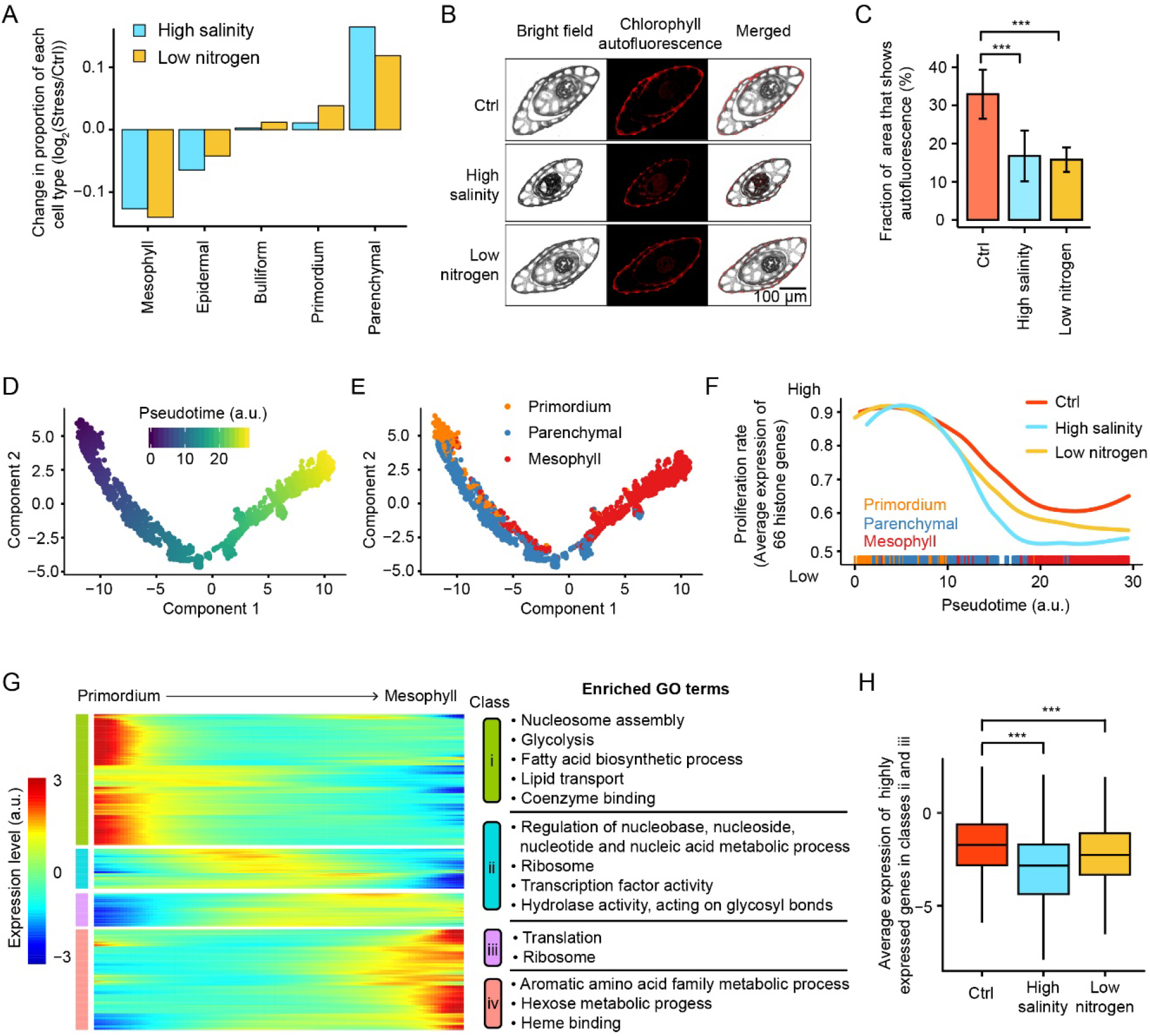
Compositional responses of cell populations upon abiotic stresses. (A) Changes in the proportion of cell types upon high-salinity or low-nitrogen stresses. The proportion was calculated by dividing the number of cells of a type by the total number of cells captured in the sample. (B–C) Microscopic images of the horizontal slices of rice seedlings grown in three conditions. Red autofluorescence is mainly from the chloroplasts in mesophyll cells. The fraction of the plant area that shows red autofluorescence was compared among three samples using the Student’s *t*-test (*N* = 10 for each treatment; ***, *P* < 0.001). (D–E) Developmental trajectory of the differentiation from primordium to parenchymal and to mesophyll cells. Each dot represents a cell and is colored by pseudotime (D) or cell types (E). (F) The proliferation rate (as inferred from the average expression levels of 66 histone genes in each cell) decreases along the developmental pseudotime, especially under abiotic stresses. We normalized the proliferation rates among three conditions according to the fast-proliferating pluripotent cells (those ranking in the first one fifth by pseudotime). (G) The expression of highly dynamic genes over the developmental trajectory. Genes are grouped into 4 classes based on the temporal expression pattern. Enriched GO terms are shown for each class. (H) Highly expressed genes in classes ii and iii are suppressed in parenchyma cells in both abiotic stress-treated samples, echoing their reduced proliferation rate in the parenchymal-mesophyll transition (F). *P* values are given by the Mann-Whitney *U* tests (***, *P* < 0.001).

To understand the underlying developmental mechanisms, we reconstructed the developmental trajectory from primordium to mesophyll and parenchymal cells. We observed a major developmental trajectory from primordium to mesophyll cells, with parenchymal cells as an intermediate (**Figures 4D–E**), echoing that mesophyll cells are mostly made up of folded parenchymal cells.

To study the effect of abiotic stresses on the proliferation and differentiation of cells we inferred the rate of cell proliferation from the average expression level of 66 histone-encoding genes, which are specifically expressed during the DNA synthesis phase. We observed a significant reduction in the proliferation rate over the developmental trajectory in all three conditions (**Figure 4F**, Spearman’s correlation coefficient *ρ* = −0.39, *P* < 10^−100^). Notably, cell proliferation was substantially suppressed during the parenchymal-mesophyll transition upon high-salinity or low-nitrogen treatment (**Figure 4F**), explaining the compositional change in cell populations (**Figure 4A**).

To further dissect the regulatory mechanisms that slow down the differentiation towards mesophyll cells upon stresses, we identified 1496 highly variable genes in the development trajectory. These genes fell into four distinct gene classes and depicted waves of gene expression in a well-organized temporal order throughout the development (**Figures 4G–H**). Genes expressed in the middle of the pseudotime axis (classes ii and iii) were associated with the transition from parenchymal to mesophyll cells. They were enriched in the GO terms related to nucleotide metabolism and translation, and their expression was suppressed in parenchymal cells upon stress treatments (**Figure 4H**), echoing the delayed development towards mesophyll cells (**Figure 4F**). Since parenchyma cells play a role in lipid storage and metabolism which are often associated with stress responses (Dionisio-Sese and Tobita, 1998; Huang, 2018), the postponed development towards mesophyll cells not only benefit rice seedlings by reducing the energy spent on photosynthesis but also by promoting the resistance to harsh environments.

## DISCUSSION

In this study, we identified all major cell types of the aerial part of rice seedlings and reconstructed their developmental trajectories using scRNA-seq. We recapitulated the transcriptional, cellular, and developmental responses to environmental stresses. Three layers of information were revealed from the single-cell transcriptome data —cell-type-specific differential gene expression, alteration in cell size, and compositional change in cell populations. Our study showcases the power of single-cell transcriptome analyses in understanding heterogeneity among plant cells.

There are some caveats in this study. First, we discarded genes whose expression may change during protoplasting to avoid artifacts associated with the preparation of scRNA-seq libraries. However, these genes may also play a role in responding to stresses. In that case, our study may miss some genes involved in the transcriptional response to stresses. Second, as a very first attempt to apply scRNA-seq to the aerial part of rice samples, replicates of the same treatment were not performed because hundreds of individual cells can serve as replicates (**Figure 2B** and **Figure S2**). Instead, two stresses (low-nitrogen and high-salinity) were treated, and shared transcriptional responses were observed (**Figure S3C** and **Figure 3E**). Third, the efficiency of capturing transcripts during the preparation of scRNA-seq libraries may be affected by abiotic stresses. If so, such effect could have led to variation in total UMI among samples, which has been used to infer cell size in this study (**Figure 3A**). Nevertheless, the size of epidermal cells was further validated with microscopy (**Figures 3B–C**). Fourth, in plants the volume of vacuole in a cell, which rarely contains mRNAs, also affects the cell size. Therefore, experimental validation with microscopy is essential for future studies when cell sizes are inferred from the total UMI captured in a plant cell. Fifth, the efficiency of protoplasting may differ among treatments. Therefore, we validated the variation in the population of mesophyll cells upon stress treatments with additional microscopic data (**Figures 4B–C** and **Figure S4**). Single-cell assay for transposase-accessible chromatin (ATAC)-seq or scRNA-seq on nuclei could partially solve this problem in the future since the extraction of nuclei is likely more uniform across treatments.

We observed variation in the level of response to stresses among cell types. More DEGs were identified in the mesophyll, parenchyma, and epidermal cells, while less were identified in primordium cells (**Figure 2C** and **Figure S3A**). This is consistent with the previous observations in heat-shock treated Arabidopsis roots that cell-type-specific expression changes often took place in differentiated cells (Jean-Baptiste et al., 2019), although we could not rule out the possibility that the detection of DEGs was statistically less sensitive due to the relatively smaller number of primordium cells. In addition, single-cell epigenetic or transcription diversity has been reported as a hallmark of developmental potential (Gulati et al., 2020; Huan et al., 2018). It would be of interest in the future to test this idea with plant single-cell transcriptome data.

In the future, scRNA-seq analyses reported in this study, together with single-cell epigenetic approaches (Buenrostro et al., 2015; Hainer et al., 2019; Huan et al., 2018; Lai et al., 2018) and methods with spatial information using *in situ* reverse transcription, can help characterize the spatiotemporal transcriptome atlas of plants. Such studies will not only promote the understanding of plant cellular and developmental biology at single-cell resolution but also help breed better crops against environmental stresses (Rhee et al., 2019).

## METHODS

### Plant materials and growth conditions

Seeds of *Oryza sativa* ssp. *japonica* variety Nipponbare were germinated on wet sterile filter paper. Seedlings were grown in a growth chamber under a 12-hour light (30°C) / 12-hour dark (28°C) photoperiod. For the control sample, seedlings were grown in the Kimura B solution for 14 days. For the high-salinity treatment, seedlings were grown in the Kimura B solution for 11 days and transferred to the high-salinity solution (Kimura B solution plus 200mM NaCl) for another three days. For the low-nitrogen treatment, seedlings were grown in the Kimura B solution for nine days and were transferred to the low-nitrogen solution (Kimura B solution only containing 20 mM NH_4_^+^) for another five days. The aerial parts of these two-week-old seedlings were harvested for scRNA-seq analyses.

### Isolation of protoplasts

Rice protoplasts were isolated from healthy two-week-old seedlings using a previously described protocol (Jabnoune et al., 2015) with modifications. Briefly, stem and sheath sections of seedlings were chopped into 0.5–1 mm strips with freshly sharpened blades. The strips were transferred immediately to freshly prepared enzyme solution, including 1.5% (w/v) cellulase R10, 0.4% (w/v) macerozyme R10, 0.6 M mannitol, 20 mM KCl, 20 mM MES (pH = 5.5), and 0.1% BSA. Plant materials (0.5 g) were incubated in 5 ml enzyme solution in dark at room temperature. After 3-hour gentle shaking, the digestion mixture was sequentially filtered through a 70 μm (FALCON, Cat # 352350) and a 40 μm (FALCON, Cat # 352340) nylon mesh sieve. Protoplasts were collected after soft washing twice (at 300 g for 3 minutes with a swinging bucket rotor) and were re-suspended in 8% mannitol solution. The protoplasts were stained by trypan blue and the activity and quantity were measured with hemocytometer counting chambers.

### Bulk RNA-seq

Total RNA was extracted from fresh plant materials with TRIzol. To detect the impact of protoplasting on the transcriptome, the total RNA was extracted before and after protoplasting, with at least five biological replications each. Strand-specific RNA-seq libraries were constructed, which were further sequenced with the Illumina sequencing platform in the paired-end 150-nt mode.

Raw reads were filtered after quality control; reads of > 50% bases with a Phred score < 20 were discarded. Adaptors were cut by fastp (Shifu et al., 2018). Clean reads were aligned to the reference Nipponbare genome (release 7 of the MSU Rice Genome Annotation Project, RGAP7) (Kawahara et al., 2013) with TopHat2 (version 2.1.1) (Kim et al., 2013). Gene expression levels were calculated from uniquely mapped reads by using HTSeq (version 0.11.2) (Anders et al., 2015).

### scRNA-seq

#### Library preparation

The scRNA-seq libraries were constructed with fresh protoplasts following the 10x Genomics Single Cell 3′ Reagent Kits v2 protocol and were sequenced with Illumina HiSeq in the paired-end 150-nt mode.

#### Estimation of the gene expression level in individual cells and identification of highly variable genes

Sequencing reads were aligned to rice reference (RGAP7) (Kawahara et al., 2013) with STAR (version 2.6.0a) (Dobin et al., 2013), and an expression matrix was generated, where each row was a feature (gene expression level) and each column was a valid cell barcode. The valid cell barcodes were determined with Cell Ranger (version 2.0.2, 10x Genomics) based on the distribution of the UMI among all barcodes (Zheng et al., 2017). The gene expression level for each cell was estimated in the unit of count per million (CPM), which controls for the total number of UMIs in a cell.

An expressed gene was defined as those captured by at least five UMIs in > 20 cells. The average and the coefficient of variation (CV) of each expressed gene were calculated among all cells. To control for the average expression level during the identification of the highly variable gene, we ordered all expressed genes by the average expression level. If the CV of a gene was greater than the median of CV among the 100 flanking genes centered at the gene by 0.2, the gene was considered as a highly variable gene. With this criterion, 1322, 855, and 1283 highly variable genes were identified in the control, high-salinity, and low-nitrogen samples, respectively. The union of these genes was used to further analyses.

#### Dimension reduction and cell clustering

With the expression matrix of the high-variable genes, the dimension reduction was perform with UMAP (R package version 0.2.2.0, under parameters n_neighbors = 5, metric = “cosine”, and min_dist = 0.01) (McInnes et al., 2018). To correct for the batch effect among samples, the mutual nearest-neighbor (MNN) method was employed by executing function fastMNN in R package Scran (version 1.10.2) (Haghverdi et al., 2018). To classify cells into clusters, function hdbscan in R package dbscan (version 1.1.3,) (McInnes et al., 2017) was performed under the parameter minPts = 40.

#### Assignment of clusters to cell types

To annotate cell clusters, we retrieved the expression level of cell-type-specific genes from a previous dataset (Jiao et al., 2009), in which marker genes were defined as genes with 1) the highest average expression in one cell type and 2) higher than three times of the average expression level in the cell type that ranked in the 2^nd^ place. In addition, cell cluster-specific genes were identified from our scRNA-seq data. They were genes with the highest average expression level in a cell cluster (among the five clusters) and 2) the highest proportion of cells expressing the gene in the cluster. In addition, these two features in the 1^st^ place cluster were required to be at least three times of the 2^nd^ place cluster and five times of the 3^rd^ place cluster. According to the biological functions of these cluster-specific genes, all five cell types were identified or validated.

#### Identification of cell-type-specific stress-response genes

DEGs for each cell type were identified as genes with at least 2-fold differences in the average expression level and *P* < 0.01 in the Mann-Whitney *U* test between a stress-treated sample and a control sample. Therefore, the sample sizes were equal to the number of cells identified in each type. The intersection of DEGs among cell types was visualized using function upset in R package UpSetR (version 1.4.0) (Lex et al., 2014).

#### Developmental trajectory

The developmental trajectories from primordium cells to other cells were performed using R package Monocle (version 2.10.1) (Satija et al., 2015). The expression levels of the 1620 high variable genes were used in the estimation of pseudotime. Genes expressed at various developmental stages were identified with function differentialGeneTest under parameter FDR < 0.1.

#### GO enrichment analyses

The GO enrichment analyses were performed on the website AgriGO2 (http://systemsbiology.cau.edu.cn/agriGOv2) (Tian et al., 2017).

### Microscopy

The aerial part of rice seedlings was cleared in ethanol:acetic acid (7:1, v/v) solution at room temperature for 14 hours as described previously (Raissig et al., 2017). The images of epidermal cells were taken with a microscope (Zeiss Axiovert 200 M) in the differential interference contrast imagining mode. Images were analyzed with ImageJ (version 1.49) (Schneider et al., 2012) and the areas of epidermal cells were estimated.

To estimate the fraction of mesophyll cells in the aerial part of rice seedlings, samples were fixed with a half-cut styrofoam board and were sliced with a freshly sharpened blade. The red autofluorescence emitted from chloroplasts was captured with a fluorescence microscope (Nikon Eclipse Ti-S). Fractions of areas showing red autofluorescence were estimated by ImageJ.

### Accession numbers

All sequencing data have been deposited to the Genome Sequence Archive (Wang et al., 2017) in BIG Data Center (http://bigd.big.ac.cn/gsa), Beijing Institute of Genomics, Chinese Academy of Sciences.

## DECLARATIONS

### Authors’ contribution

Q.H., K.L., and W.Q. designed the experiments; Q.H., Y.W., and K.L. performed the experiments; Y.W., X.C., and Q.H. analyzed the data; Q.H., Y.W., and W.Q. wrote the manuscript.

## Acknowledgments

We thank Ying Chen for support in data analysis, and Caihuan Tian and Taolan Zhao for discussion.

## REFERENCES

Anders, S., Pyl, P.T., and Huber, W. (2015). HTSeq--a Python framework to work with high-throughput sequencing data. Bioinformatics 31, 166–169.

Birnbaum, K., Shasha, D.E., Wang, J.Y., Jung, J.W., Lambert, G.M., Galbraith, D.W., and Benfey, P.N. (2003). A gene expression map of the Arabidopsis root. Science 302, 1956–1960.

Buenrostro, J.D., Wu, B., Litzenburger, U.M., Ruff, D., Gonzales, M.L., Snyder, M.P., Chang, H.Y., and Greenleaf, W.J. (2015). Single-cell chromatin accessibility reveals principles of regulatory variation. Nature 523, 486–490.

Chen, Y., Li, K., Chu, X., Carey, L.B., and Qian, W. (2019). Synchronized replication of genes encoding the same protein complex in fast-proliferating cells. Genome Res 29, 1929–1938.

Denyer, T., Ma, X., Klesen, S., Scacchi, E., Nieselt, K., and Timmermans, M.C.P. (2019). Spatiotemporal Developmental Trajectories in the Arabidopsis Root Revealed Using High-Throughput Single-Cell RNA Sequencing. Dev Cell 48, 840–852 e845.

Dinneny, J.R., Long, T.A., Wang, J.Y., Jung, J.W., Mace, D., Pointer, S., Barron, C., Brady, S.M., Schiefelbein, J., and Benfey, P.N. (2008). Cell identity mediates the response of Arabidopsis roots to abiotic stress. Science 320, 942–945.

Dionisio-Sese, M.L., and Tobita, S. (1998). Antioxidant responses of rice seedlings to salinity stress. Plant Science 135, 1–9.

Dobin, A., Davis, C.A., Schlesinger, F., Drenkow, J., Zaleski, C., Jha, S., Batut, P., Chaisson, M., and Gingeras, T.R. (2013). STAR: ultrafast universal RNA-seq aligner. Bioinformatics 29, 15–21.

Efroni, I., Ip, P.L., Nawy, T., Mello, A., and Birnbaum, K.D. (2015). Quantification of cell identity from single-cell gene expression profiles. Genome Biol 16, 9.

Frank, M.H., and Scanlon, M.J. (2015). Cell-specific transcriptomic analyses of three-dimensional shoot development in the moss Physcomitrella patens. Plant J 83, 743–751.

Gulati, G.S., Sikandar, S.S., Wesche, D.J., Manjunath, A., Bharadwaj, A., Berger, M.J., Ilagan, F., Kuo, A.H., Hsieh, R.W., Cai, S., et al. (2020). Single-cell transcriptional diversity is a hallmark of developmental potential. Science 367, 405–411.

Haghverdi, L., Lun, A.T.L., Morgan, M.D., and Marioni, J.C. (2018). Batch effects in single-cell RNA-sequencing data are corrected by matching mutual nearest neighbors. Nat Biotechnol 36, 421–427.

Hainer, S.J., Boskovic, A., McCannell, K.N., Rando, O.J., and Fazzio, T.G. (2019). Profiling of Pluripotency Factors in Single Cells and Early Embryos. Cell 177, 1319–1329 e1311.

Han, Y., Chu, X., Yu, H., Ma, Y.-K., Wang, X.-J., Qian, W., and Jiao, Y. (2017). Single-cell transcriptome analysis reveals widespread monoallelic gene expression in individual rice mesophyll cells. Science Bulletin 62, 1304–1314.

Huan, Q., Zhang, Y., Wu, S., and Qian, W. (2018). HeteroMeth: A Database of Cell-to-cell Heterogeneity in DNA Methylation. Genomics Proteomics Bioinformatics 16, 234–243.

Huang, A.H.C. (2018). Plant Lipid Droplets and Their Associated Proteins: Potential for Rapid Advances. Plant Physiol 176, 1894–1918.

Jabnoune, M., Secco, D., Lecampion, C., Robaglia, C., Shu, Q., and Poirier, Y. (2015). An Efficient Procedure for Protoplast Isolation from Mesophyll Cells and Nuclear Fractionation in Rice. BIO-PROTOCOL 5.

Jean-Baptiste, K., McFaline-Figueroa, J.L., Alexandre, C.M., Dorrity, M.W., Saunders, L., Bubb, K.L., Trapnell, C., Fields, S., Queitsch, C., and Cuperus, J.T. (2019). Dynamics of Gene Expression in Single Root Cells of Arabidopsis thaliana. Plant Cell 31, 993–1011.

Jiao, Y., Tausta, S.L., Gandotra, N., Sun, N., Liu, T., Clay, N.K., Ceserani, T., Chen, M., Ma, L., Holford, M., et al. (2009). A transcriptome atlas of rice cell types uncovers cellular, functional and developmental hierarchies. Nat Genet 41, 258–263.

Kawahara, Y., de la Bastide, M., Hamilton, J.P., Kanamori, H., McCombie, W.R., Ouyang, S., Schwartz, D.C., Tanaka, T., Wu, J., Zhou, S., et al. (2013). Improvement of the Oryza sativa Nipponbare reference genome using next generation sequence and optical map data. Rice (N Y) 6, 4.

Kempe, H., Schwabe, A., Cremazy, F., Verschure, P.J., and Bruggeman, F.J. (2015). The volumes and transcript counts of single cells reveal concentration homeostasis and capture biological noise. Mol Biol Cell 26, 797–804.

Kim, D., Pertea, G., Trapnell, C., Pimentel, H., Kelley, R., and Salzberg, S.L. (2013). TopHat2: accurate alignment of transcriptomes in the presence of insertions, deletions and gene fusions. Genome Biol 14, R36.

Klein, A.M., Mazutis, L., Akartuna, I., Tallapragada, N., Veres, A., Li, V., Peshkin, L., Weitz, D.A., and Kirschner, M.W. (2015). Droplet barcoding for single-cell transcriptomics applied to embryonic stem cells. Cell 161, 1187–1201.

Lai, B., Gao, W., Cui, K., Xie, W., Tang, Q., Jin, W., Hu, G., Ni, B., and Zhao, K. (2018). Principles of nucleosome organization revealed by single-cell micrococcal nuclease sequencing. Nature 562, 281–285.

Lex, A., Gehlenborg, N., Strobelt, H., Vuillemot, R., and Pfister, H. (2014). UpSet: Visualization of Intersecting Sets. IEEE Transactions on Visualization and Computer Graphics 20, 1983–1992.

Lieckfeldt, E., Simon-Rosin, U., Kose, F., Zoeller, D., Schliep, M., and Fisahn, J. (2008). Gene expression profiling of single epidermal, basal and trichome cells of Arabidopsis thaliana. J Plant Physiol 165, 1530–1544.

Macosko, E.Z., Basu, A., Satija, R., Nemesh, J., Shekhar, K., Goldman, M., Tirosh, I., Bialas, A.R., Kamitaki, N., Martersteck, E.M., et al. (2015). Highly Parallel Genome-wide Expression Profiling of Individual Cells Using Nanoliter Droplets. Cell 161, 1202–1214.

McInnes, L., Healy, J., and Astels, S. (2017). hdbscan: Hierarchical density based clustering. Journal of Open Source Software 2, 205.

McInnes, L., Healy, J., and Melville, J. (2018). UMAP: Uniform Manifold Approximation and Projection for Dimension Reduction. arXiv:1802.03426.

Raissig, M.T., Matos, J.L., Anleu Gil, M.X., Kornfeld, A., Bettadapur, A., Abrash, E., Allison, H.R., Badgley, G., Vogel, J.P., Berry, J.A., et al. (2017). Mobile MUTE specifies subsidiary cells to build physiologically improved grass stomata. Science 355, 1215–1218.

Rhee, S.Y., Birnbaum, K.D., and Ehrhardt, D.W. (2019). Towards Building a Plant Cell Atlas. Trends Plant Sci 24, 303–310.

Rich-Griffin, C., Stechemesser, A., Finch, J., Lucas, E., Ott, S., and Schafer, P. (2019). Single-Cell Transcriptomics: A High-Resolution Avenue for Plant Functional Genomics. Trends Plant Sci.

Ryu, K.H., Huang, L., Kang, H.M., and Schiefelbein, J. (2019). Single-Cell RNA Sequencing Resolves Molecular Relationships Among Individual Plant Cells. Plant Physiol 179, 1444–1456.

Satija, R., Farrell, J.A., Gennert, D., Schier, A.F., and Regev, A. (2015). Spatial reconstruction of single-cell gene expression data. Nat Biotechnol 33, 495–502.

Schneider, C.A., Rasband, W.S., and Eliceiri, K.W. (2012). NIH Image to ImageJ: 25 years of image analysis. Nat Methods 9, 671–675.

Shulse, C.N., Cole, B.J., Ciobanu, D., Lin, J., Yoshinaga, Y., Gouran, M., Turco, G.M., Zhu, Y., O’Malley, R.C., Brady, S.M., et al. (2019). High-Throughput Single-Cell Transcriptome Profiling of Plant Cell Types. Cell Rep 27, 2241–2247 e2244.

Tian, T., Liu, Y., Yan, H., You, Q., Yi, X., Du, Z., Xu, W., and Su, Z. (2017). agriGO v2.0: a GO analysis toolkit for the agricultural community, 2017 update. Nucleic Acids Res 45, W122–W129.

Wang, Y., Song, F., Zhu, J., Zhang, S., Yang, Y., Chen, T., Tang, B., Dong, L., Ding, N., Zhang, Q., et al. (2017). GSA: Genome Sequence Archive. Genomics Proteomics Bioinformatics 15, 14–18.

Zhang, T.Q., Xu, Z.G., Shang, G.D., and Wang, J.W. (2019). A Single-Cell RNA Sequencing Profiles the Developmental Landscape of Arabidopsis Root. Mol Plant 12, 648–660.

Zheng, G.X., Terry, J.M., Belgrader, P., Ryvkin, P., Bent, Z.W., Wilson, R., Ziraldo, S.B., Wheeler, T.D., McDermott, G.P., Zhu, J., et al. (2017). Massively parallel digital transcriptional profiling of single cells. Nat Commun 8, 14049.

