## Supplementary information for "Single-cell transcriptome analyses recapitulate the cellular and developmental responses to abiotic stresses in rice"

Supplementary Information includes Supplementary Figures S1-4 and Supplementary Tables S1-3.

**SUPPLEMENTARY FIGURES**

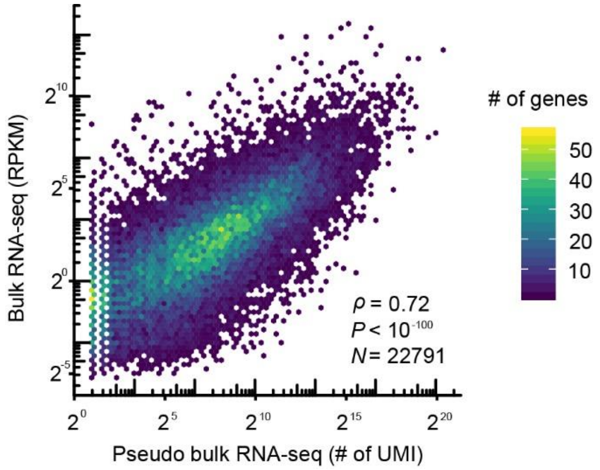


**Figure S1. Comparison of gene expression between scRNA-seq and bulk RNA-seq libraries.**

All scRNA-seq reads were aggregated from individual cells and combined into one pseudobulk profile. Spearman's correlation coefficient *ρ*, the corresponding *P* value, and the number of genes (*N*) are shown. The color indicates dot density.

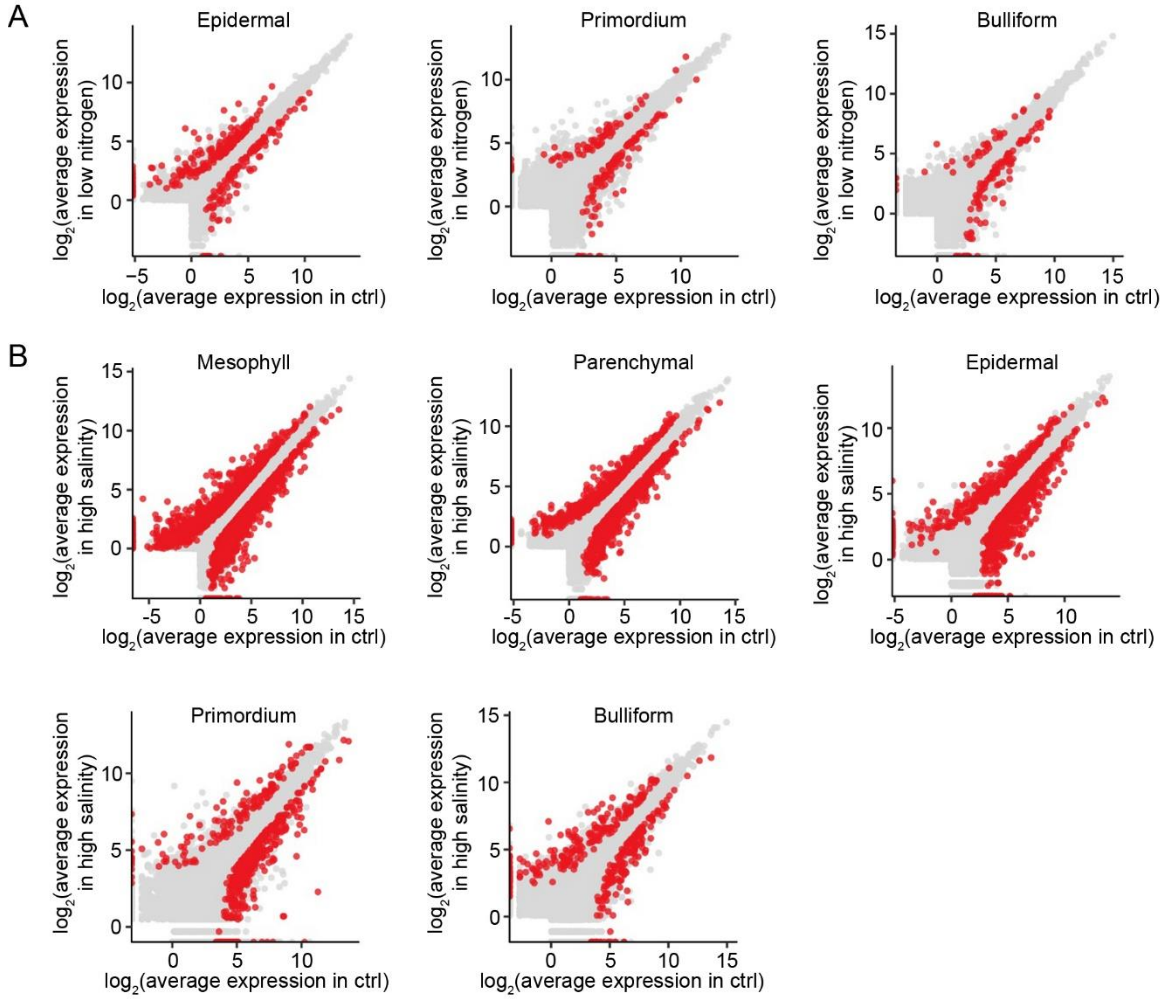


**Figure S2. DEGs identified upon abiotic stresses in individual cell types.** Similar to Figure 2B.

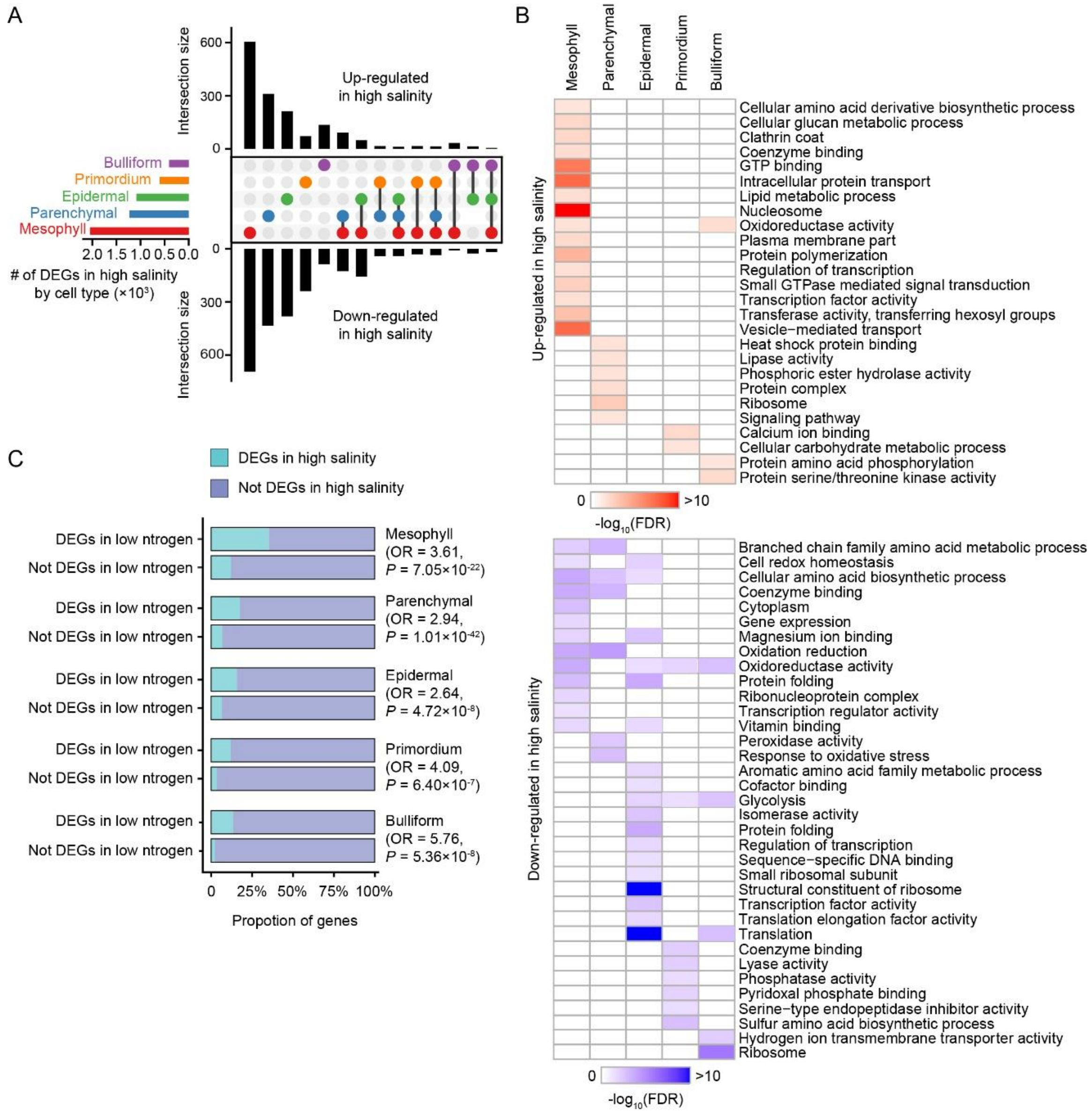


**Figure S3. Transcriptome responses upon the high-salinity and low-nitrogen stresses in individual cell types.**

(A–B) Similar to Figure 2C-D, for the high-salinity stress.

(C) DEGs are significantly overlapped between two abiotic stress-treated samples in all five cell types. Odds ratios and *P* values are given by Fisher's exact tests.

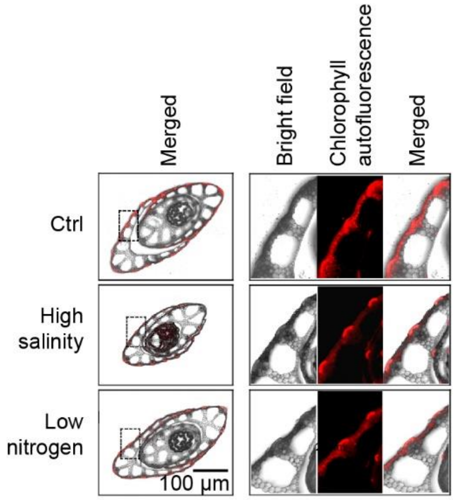


**Figure S4. The fraction of plant area that shows chlorophyll autofluorescence.** The dashed area of the left penal (the same as Figure 4B) is zoomed in on the right.

### SUPPLEMENTARY TABLES

**Table S1. Summary of the scRNA-seq data.**

| Terms | Ctrl | High salinity | Low nitrogen |
|---|---|---|---|
| Estimated number of cells | 2027 | 783 | 1905 |
| Median total UMI per cell | 11966 | 8959 | 9392 |
| Median number of genes per cell | 2678 | 2178 | 2410 |
| Reads mapped confidently to transcriptome | 75.90% | 75.20% | 74.80% |
| Total genes detected | 25263 | 22513 | 25390 |
| Non-TE related genes detected | 23802 | 21323 | 23886 |

**Table S2. DEGs during protoplasting.**

| Locus | Fold change* | FDR** | Annotation |
|---|---|---|---|
| LOC_Os01g08670 | 12.956 | 0.0085 | expressed protein |
| LOC_Os01g13420 | 0.078 | 0.0026 | SOUL heme-binding protein, putative, expressed |
| LOC_Os01g15120 | 0.071 | 0.0035 | hydrolase, alpha/beta fold family domain containing protein, expressed |
| LOC_Os01g16180 | 0.059 | 0.0085 | MT-A70 domain containing protein, expressed |
| LOC_Os01g25700 | 0.052 | 0.0071 | transposon protein, putative, CACTA, En/Spm sub-class, expressed |
| LOC_Os01g37690 | 0.029 | 0.0020 | sodium/calcium exchanger protein, putative, expressed |
| LOC_Os01g43230 | 0.010 | 0.0005 | expressed protein |
| LOC_Os01g64010 | 0.181 | 0.0035 | dual specificity protein phosphatase, putative, expressed |
| LOC_Os01g68870 | 0.122 | 0.0005 | leucine-rich repeat receptor protein kinase EXS precursor, putative, expressed |
| LOC_Os01g70400 | 0.276 | 0.0020 | expressed protein |
| LOC_Os02g06300 | 0.109 | 0.0014 | GTP-binding protein lepA, putative, expressed |
| LOC_Os02g13510 | 0.012 | 0.0013 | receptor-like protein kinase 5 precursor, putative, expressed |
| LOC_Os02g14280 | 0.060 | 0.0080 | transposon protein, putative, unclassified, expressed |
| LOC_Os02g20410 | 0.162 | 0.0071 | GTP binding protein, putative, expressed |

| | | | |
|---|---|---|---|
| LOC_Os02g27060 | 0.061 | 0.0000 | high mobility group, putative, expressed |
| LOC_Os02g30140 | 0.088 | 0.0029 | expressed protein |
| LOC_Os02g30530 | 0.216 | 0.0071 | transposon protein, putative, unclassified, expressed |
| LOC_Os02g45054 | 0.112 | 0.0031 | ZOS2-15 - C2H2 zinc finger protein, expressed |
| LOC_Os02g51760 | 0.135 | 0.0084 | expressed protein |
| LOC_Os02g56510 | 0.029 | 0.0076 | phosphate transporter 1, putative, expressed |
| LOC_Os02g57750 | 0.160 | 0.0085 | protein binding protein, putative, expressed |
| LOC_Os02g58150 | 0.017 | 0.0048 | expressed protein |
| LOC_Os03g24760 | 0.150 | 0.0004 | terpene synthase, putative, expressed |
| LOC_Os03g29190 | 0.050 | 0.0021 | PDI, putative, expressed |
| LOC_Os03g45280 | 15.000 | 0.0013 | dehydrin, putative, expressed |
| LOC_Os03g59340 | 0.057 | 0.0035 | CESA2 - cellulose synthase, expressed |
| LOC_Os04g04060 | 0.029 | 0.0031 | dynamin family protein, putative, expressed |
| LOC_Os04g20474 | 0.016 | 0.0020 | UDP-glucoronosyl and UDP-glucosyl transferase domain containing protein, expressed |
| LOC_Os04g34610 | 0.036 | 0.0071 | expressed protein |
| LOC_Os04g36062 | 0.046 | 0.0085 | expressed protein |
| LOC_Os04g39840 | 0.021 | 0.0005 | Os4bglu10 - beta-glucosidase homologue, similar to Os4Bglu12 exoglucanase/beta-glucosidase, expressed |

| | | | |
|---|---|---|---|
| LOC_Os04g45470 | 0.160 | 0.0020 | vacuolar-processing enzyme precursor, putative, expressed |
| LOC_Os04g49690 | 5.165 | 0.0035 | FERONIA receptor-like kinase, putative, expressed |
| LOC_Os04g52710 | 0.079 | 0.0080 | imidazoleglycerol-phosphate dehydratase, putative, expressed |
| LOC_Os04g57500 | 0.132 | 0.0020 | phosphatidate cytidylyltransferase, putative, expressed |
| LOC_Os05g03490 | 0.104 | 0.0005 | expressed protein |
| LOC_Os05g04180 | 0.157 | 0.0080 | agenet domain containing protein, putative, expressed |
| LOC_Os05g09380 | 0.051 | 0.0033 | AR791, putative, expressed |
| LOC_Os05g15140 | 0.121 | 0.0035 | expressed protein |
| LOC_Os05g33620 | 0.096 | 0.0037 | expressed protein |
| LOC_Os05g36160 | 0.103 | 0.0057 | G-box-binding factor 4, putative, expressed |
| LOC_Os05g50990 | 0.042 | 0.0016 | TTL3, putative, expressed |
| LOC_Os06g05470 | 264.367 | 0.0008 | expressed protein |
| LOC_Os06g05630 | 0.017 | 0.0071 | GDSL-like lipase/acylhydrolase, putative, expressed |
| LOC_Os06g17950 | 0.088 | 0.0005 | NBS-LRR disease resistance protein, putative, expressed |
| LOC_Os07g06080 | 0.095 | 0.0080 | FAD dependent oxidoreductase domain containing protein, expressed |
| LOC_Os07g10630 | 0.041 | 0.0013 | expressed protein |
| LOC_Os08g01490 | 0.058 | 0.0003 | cytochrome P450, putative, expressed |
| LOC_Os08g29170 | 0.227 | 0.0011 | dehydrogenase, putative, expressed |

| | | | |
|---|---|---|---|
| LOC_Os08g39370 | 0.057 | 0.0013 | citrate transporter, putative, expressed |
| LOC_Os09g07360 | 0.035 | 0.0014 | frigida, putative, expressed |
| LOC_Os09g19570 | 0.052 | 0.0054 | AGAP009532-PA, putative, expressed |
| LOC_Os09g34980 | 0.232 | 0.0037 | zinc knuckle family protein, expressed |
| LOC_Os09g36240 | 0.034 | 0.0080 | deoxyribodipyrimidine photolyase family protein, expressed |
| LOC_Os09g36360 | 0.124 | 0.0092 | protein of unknown function containing protein, expressed |
| LOC_Os10g36000 | 0.156 | 0.0041 | remorin C-terminal domain containing protein, putative, expressed |
| LOC_Os10g36260 | 0.241 | 0.0038 | expressed protein |
| LOC_Os10g41780 | 0.109 | 0.0013 | chlorophyllide a oxygenase, chloroplast precursor, putative, expressed |
| LOC_Os11g09850 | 0.141 | 0.0071 | expressed protein |
| LOC_Os11g17970 | 0.103 | 0.0085 | POT family protein, expressed |
| LOC_Os11g42030 | 0.128 | 0.0031 | expressed protein |
| LOC_Os12g03270 | 0.177 | 0.0093 | ELMO/CED-12 family protein, putative, expressed |
| LOC_Os12g04480 | 0.087 | 0.0054 | cytochrome P450, putative, expressed |

* Fold change was calculated as the ratio between the average expression levels after and before protoplasting in the bulk RNA-seq samples.

** FDR was corrected from the *P* values from linear regression models.

**Table S3. List of cluster-specific genes.**

| Locus | Cluster | Annotation |
|---|---|---|
| LOC_Os01g58049 | 1 | photosystem I assembly protein ycf4, putative, expressed |
| LOC_Os02g24642 | 1 | photosystem II reaction center protein K precursor, putative, expressed |
| LOC_Os03g01300 | 1 | LTPL114 - Protease inhibitor/seed storage/LTP family protein precursor, expressed |
| LOC_Os04g16866 | 1 | photosystem II reaction center protein K precursor, putative, expressed |
| LOC_Os04g24410 | 1 | kinesin heavy chain isolog, putative, expressed |
| LOC_Os05g30760 | 1 | esterase, putative, expressed |
| LOC_Os08g15296 | 1 | photosystem II reaction center protein H, putative, expressed |
| LOC_Os08g39840 | 1 | lipoxygenase, chloroplast precursor, putative, expressed |
| LOC_Os10g21268 | 1 | ribulose bisphosphate carboxylase large chain precursor, putative, expressed |
| LOC_Os10g35950 | 1 | transferase family protein, putative, expressed |
| LOC_Os01g68589 | 2 | LTPL39 - Protease inhibitor/seed storage/LTP family protein precursor, expressed |
| LOC_Os04g52260 | 2 | LTPL124 - Protease inhibitor/seed storage/LTP family protein precursor, expressed |
| LOC_Os04g55159 | 2 | LTPL125 - Protease inhibitor/seed storage/LTP family protein precursor, putative, expressed |
| LOC_Os07g46480 | 2 | eukaryotic aspartyl protease domain containing protein, expressed |
| LOC_Os08g35760 | 2 | Cupin domain containing protein, expressed |
| LOC_Os11g25780 | 2 | PB1 domain containing protein, expressed |

| | | |
|---|---|---|
| LOC_Os01g73170 | 3 | peroxidase precursor, putative, expressed |
| LOC_Os02g09240 | 3 | cytochrome P450 71D8, putative, expressed |
| LOC_Os03g17150 | 3 | ZOS3-09 - C2H2 zinc finger protein, expressed |
| LOC_Os04g01140 | 3 | cytochrome P450 93A2, putative, expressed |
| LOC_Os04g39350 | 3 | heavy metal associated domain containing protein, expressed |
| LOC_Os05g19000 | 3 | expressed protein |
| LOC_Os05g31750 | 3 | annexin, putative, expressed |
| LOC_Os05g40384 | 3 | cytochrome P450, putative, expressed |
| LOC_Os05g51240 | 3 | hydrolase, alpha/beta fold family domain containing protein, expressed |
| LOC_Os06g20920 | 3 | SAM dependent carboxyl methyltransferase, putative, expressed |
| LOC_Os06g42680 | 3 | phytosulfokines 1 precursor, putative, expressed |
| LOC_Os06g49880 | 3 | B-box zinc finger family protein, putative, expressed |
| LOC_Os07g46870 | 3 | sex determination protein tasselseed-2, putative, expressed |
| LOC_Os08g02230 | 3 | FAD-binding and arabino-lactone oxidase domains containing protein, putative, expressed |
| LOC_Os08g30020 | 3 | membrane protein, putative, expressed |
| LOC_Os08g44270 | 3 | vignain precursor, putative, expressed |
| LOC_Os09g21180 | 3 | homeobox associated leucine zipper, putative, expressed |
| LOC_Os09g28210 | 3 | bHelix-loop-helix transcription factor, putative, expressed |
| LOC_Os09g29200 | 3 | glutathione S-transferase, putative, expressed |
| LOC_Os10g34920 | 3 | secretory protein, putative, expressed |

| | | |
|---|---|---|
| LOC_Os10g38700 | 3 | glutathione S-transferase, putative, expressed |
| LOC_Os01g01650 | 4 | isoflavone reductase homolog IRL, putative, expressed |
| LOC_Os01g01660 | 4 | isoflavone reductase, putative, expressed |
| LOC_Os01g02010 | 4 | expressed protein |
| LOC_Os01g09880 | 4 | harpin-induced protein 1 domain containing protein, expressed |
| LOC_Os01g14850 | 4 | MFS18 protein precursor, putative, expressed |
| LOC_Os01g16030 | 4 | ADP-ribosylation factor, putative, expressed |
| LOC_Os01g19820 | 4 | universal stress protein domain containing protein, putative, expressed |
| LOC_Os01g47780 | 4 | fasciclin domain containing protein, expressed |
| LOC_Os01g53240 | 4 | BURP domain containing protein, expressed |
| LOC_Os01g60740 | 4 | LTPL16 - Protease inhibitor/seed storage/LTP family protein precursor, expressed |
| LOC_Os02g33550 | 4 | harpin-induced protein 1 domain containing protein, expressed |
| LOC_Os02g43194 | 4 | aldehyde dehydrogenase, putative, expressed |
| LOC_Os02g48140 | 4 | hsp20/alpha crystallin family protein, putative, expressed |
| LOC_Os02g56860 | 4 | 3-ketoacyl-CoA synthase, putative, expressed |
| LOC_Os03g08360 | 4 | 3-ketoacyl-CoA synthase 10, putative, expressed |
| LOC_Os03g14130 | 4 | POEI1 - Pollen Ole e I allergen and extensin family protein precursor, expressed |
| LOC_Os03g21040 | 4 | stress responsive protein, putative, expressed |
| LOC_Os03g45619 | 4 | cytochrome P450, putative, expressed |
| LOC_Os03g57200 | 4 | glutathione S-transferase, putative, expressed |

| | | |
|---|---|---|
| LOC_Os03g57460 | 4 | fasciclin domain containing protein, expressed |
| LOC_Os04g16450 | 4 | aquaporin protein, putative, expressed |
| LOC_Os04g21320 | 4 | membrane associated DUF588 domain containing protein, putative, expressed |
| LOC_Os04g36750 | 4 | hsp20/alpha crystallin family protein, putative, expressed |
| LOC_Os04g39150 | 4 | pathogenesis-related Bet v I family protein, putative, expressed |
| LOC_Os04g55250 | 4 | expressed protein |
| LOC_Os05g08554 | 4 | expressed protein |
| LOC_Os05g09704 | 4 | HAD superfamily phosphatase, putative, expressed |
| LOC_Os05g10210 | 4 | HAD superfamily phosphatase, putative, expressed |
| LOC_Os05g13940 | 4 | retrotransposon protein, putative, unclassified, expressed |
| LOC_Os05g38230 | 4 | oxidoreductase, aldo/keto reductase family protein, putative, expressed |
| LOC_Os05g45020 | 4 | zinc finger/CCCH transcription factor, putative, expressed |
| LOC_Os05g48900 | 4 | fasciclin domain containing protein, expressed |
| LOC_Os06g04990 | 4 | early nodulin 93 ENOD93 protein, putative, expressed |
| LOC_Os06g05010 | 4 | early nodulin 93 ENOD93 protein, putative, expressed |
| LOC_Os06g18670 | 4 | anthocyanidin 3-O-glucosyltransferase, putative, expressed |
| LOC_Os06g35970 | 4 | meiosis 5, putative, expressed |
| LOC_Os07g05950 | 4 | expressed protein |
| LOC_Os07g10440 | 4 | expressed protein |
| LOC_Os07g41360 | 4 | alpha-1,4-glucan-protein synthase, putative, expressed |

| | | |
|---|---|---|
| LOC_Os07g45060 | 4 | uncharacterized GPI-anchored protein At5g19240 precursor, putative, expressed |
| LOC_Os08g44360 | 4 | male sterility protein 2, putative, expressed |
| LOC_Os09g04710 | 4 | GDSL-like lipase/acylhydrolase, putative, expressed |
| LOC_Os09g31000 | 4 | EF hand family protein, expressed |
| LOC_Os10g05820 | 4 | POEI5 - Pollen Ole e I allergen and extensin family protein precursor, expressed |
| LOC_Os10g05860 | 4 | POEI7 - Pollen Ole e I allergen and extensin family protein precursor, putative, expressed |
| LOC_Os10g05930 | 4 | POEI10 - Pollen Ole e I allergen and extensin family protein precursor, expressed |
| LOC_Os10g05970 | 4 | POEI12 - Pollen Ole e I allergen and extensin family protein precursor, expressed |
| LOC_Os10g05980 | 4 | POEI13 - Pollen Ole e I allergen and extensin family protein precursor, expressed |
| LOC_Os10g05990 | 4 | POEI14 - Pollen Ole e I allergen and extensin family protein precursor, expressed |
| LOC_Os10g06000 | 4 | POEI15 - Pollen Ole e I allergen and extensin family protein precursor, expressed |
| LOC_Os10g30150 | 4 | universal stress protein domain containing protein, putative, expressed |
| LOC_Os10g37400 | 4 | DUF538 domain containing protein, putative, expressed |
| LOC_Os11g02080 | 4 | expressed protein |
| LOC_Os11g32890 | 4 | expressed protein |
| LOC_Os12g37650 | 4 | DUF538 domain containing protein, putative, expressed |
| LOC_Os01g06220 | 5 | gibberellin receptor GID1L2, putative, expressed |
| LOC_Os01g06500 | 5 | PHLOEM 2-LIKE A5, putative, expressed |

| | | |
|---|---|---|
| LOC_Os01g27140 | 5 | OsGrx_C7 - glutaredoxin subgroup III, expressed |
| LOC_Os01g57004 | 5 | adhesive/proline-rich protein, putative, expressed |
| LOC_Os01g70500 | 5 | expressed protein |
| LOC_Os01g70710 | 5 | heavy metal-associated domain containing protein, expressed |
| LOC_Os01g73940 | 5 | expressed protein |
| LOC_Os02g09830 | 5 | bZIP transcription factor domain containing protein, expressed |
| LOC_Os02g11020 | 5 | cytochrome P450 72A1, putative, expressed |
| LOC_Os02g45200 | 5 | dof zinc finger domain containing protein, putative, expressed |
| LOC_Os02g52540 | 5 | EF hand family protein, putative, expressed |
| LOC_Os02g53160 | 5 | tyrosine phosphatase family protein, putative, expressed |
| LOC_Os02g53180 | 5 | 1-aminocyclopropane-1-carboxylate oxidase protein, putative, expressed |
| LOC_Os03g06790 | 5 | hypothetical protein |
| LOC_Os03g09230 | 5 | LTPL69 - Protease inhibitor/seed storage/LTP family protein precursor, expressed |
| LOC_Os03g14150 | 5 | expressed protein |
| LOC_Os03g20290 | 5 | aspartic proteinase nepenthesin-1 precursor, putative, expressed |
| LOC_Os03g50960 | 5 | LTPL118 - Protease inhibitor/seed storage/LTP family protein precursor, expressed |
| LOC_Os03g58500 | 5 | expressed protein |
| LOC_Os03g60850 | 5 | peptide transporter PTR2, putative, expressed |
| LOC_Os03g61829 | 5 | haloacid dehalogenase-like hydrolase family protein, putative, expressed |

| | | |
|---|---|---|
| LOC_Os03g63390 | 5 | plastocyanin-like domain containing protein, putative, expressed |
| LOC_Os04g08350 | 5 | cysteine synthase, chloroplast/chromoplast precursor, putative, expressed |
| LOC_Os04g12720 | 5 | indole-3-acetate beta-glucosyltransferase, putative, expressed |
| LOC_Os04g50940 | 5 | peptide transporter PTR2, putative, expressed |
| LOC_Os04g56900 | 5 | transferase family protein, putative, expressed |
| LOC_Os05g03934 | 5 | expressed protein |
| LOC_Os05g04000 | 5 | expressed protein |
| LOC_Os05g05080 | 5 | expressed protein |
| LOC_Os05g05100 | 5 | expressed protein |
| LOC_Os05g05110 | 5 | expressed protein |
| LOC_Os05g06390 | 5 | retrotransposon protein, putative, unclassified, expressed |
| LOC_Os05g11600 | 5 | expressed protein |
| LOC_Os05g11610 | 5 | expressed protein |
| LOC_Os05g50770 | 5 | C4-dicarboxylate transporter/malic acid transport protein, expressed |
| LOC_Os06g07230 | 5 | tyrosine protein kinase domain containing protein, putative, expressed |
| LOC_Os06g08640 | 5 | transferase family protein, putative, expressed |
| LOC_Os06g10610 | 5 | expressed protein |
| LOC_Os06g11310 | 5 | plastocyanin-like domain containing protein, putative, expressed |
| LOC_Os06g47640 | 5 | OsCML29 - Calmodulin-related calcium sensor protein, expressed |
| LOC_Os07g03319 | 5 | SCP-like extracellular protein, expressed |

| | | |
|---|---|---|
| LOC_Os07g03580 | 5 | SCP-like extracellular protein, expressed |
| LOC_Os07g03590 | 5 | SCP-like extracellular protein, expressed |
| LOC_Os07g03600 | 5 | SCP-like extracellular protein, expressed |
| LOC_Os07g31390 | 5 | beta-expansin precursor, putative, expressed |
| LOC_Os07g35810 | 5 | TKL_IRAK_DUF26-ld.6 - DUF26 kinases have homology to DUF26 containing loci, expressed |
| LOC_Os07g37950 | 5 | expressed protein |
| LOC_Os07g43770 | 5 | expressed protein |
| LOC_Os08g04130 | 5 | copine-6, putative, expressed |
| LOC_Os08g24200 | 5 | expressed protein |
| LOC_Os08g31140 | 5 | heavy metal-associated domain containing protein, expressed |
| LOC_Os08g32970 | 5 | annexin, putative, expressed |
| LOC_Os08g41290 | 5 | AIR12, putative, expressed |
| LOC_Os09g20000 | 5 | heavy metal-associated domain containing protein, expressed |
| LOC_Os09g30414 | 5 | aspartic proteinase nepenthesin-2 precursor, putative, expressed |
| LOC_Os09g33640 | 5 | expressed protein |
| LOC_Os09g38670 | 5 | thioredoxin, putative, expressed |
| LOC_Os10g06630 | 5 | peptidyl-prolyl cis-trans isomerase, putative, expressed |
| LOC_Os10g31440 | 5 | retrotransposon protein, putative, unclassified, expressed |
| LOC_Os10g31460 | 5 | retrotransposon protein, putative, unclassified, expressed |
| LOC_Os10g34730 | 5 | GEM, putative, expressed |

| | | |
|---|---|---|
| LOC_Os10g38040 | 5 | lysM domain containing protein, putative, expressed |
| LOC_Os11g08380 | 5 | 1-aminocyclopropane-1-carboxylate oxidase, putative, expressed |
| LOC_Os11g43520 | 5 | OsGrx_C17 - glutaredoxin subgroup III, expressed |
| LOC_Os11g47580 | 5 | glycosyl hydrolase, putative, expressed |
| LOC_Os11g47680 | 5 | thaumatin family domain containing protein, expressed |
